# Mass Spectrometry-Based Plasma Proteomics: Considerations from Sample Collection to Achieving Translational Data

**DOI:** 10.1101/716563

**Authors:** Vera Ignjatovic, Philipp E Geyer, Krishnan K Palaniappan, Jessica E Chaaban, Gilbert S Omenn, Mark S Baker, Eric W Deutsch, Jochen M Schwenk

## Abstract

The proteomic analyses of human blood and blood-derived products (e.g. plasma) offers an attractive avenue to translate research progress from the laboratory into the clinic. However, due to its unique protein composition, performing proteomics assays with plasma is challenging. Plasma proteomics has regained interest due to recent technological advances, but challenges imposed by both complications inherent to studying human biology (e.g. inter-individual variability), analysis of biospecimen (e.g. sample variability), as well as technological limitations remain. As part of the Human Proteome Project (HPP), the Human Plasma Proteome Project (HPPP) brings together key aspects of the plasma proteomics pipeline. Here, we provide considerations and recommendations concerning study design, plasma collection, quality metrics, plasma processing workflows, mass spectrometry (MS) data acquisition, data processing and bioinformatic analysis. With exciting opportunities in studying human health and disease though this plasma proteomics pipeline, a more informed analysis of human plasma will accelerate interest whilst enhancing possibilities for the incorporation of proteomics-scaled assays into clinical practice.

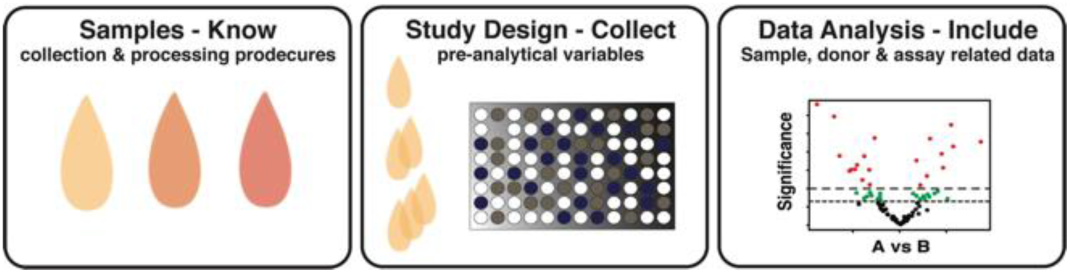

## Introduction

Blood plays a central role in facilitating diverse biological processes. Whole blood is an easily accessible and minimally invasive tissue that affords a significant opportunity to learn about human biology. Recent interests in using whole blood as a form of “liquid biopsy” for personalized medicine applications, including more effective monitoring of therapeutic response to treatment are driving the discovery of novel disease-specific biomarkers ^1,2^. Blood is a complex mixture of cells, exosomes, nucleic acids, proteins, lipids and metabolites, amongst other components. The liquid component of whole blood, termed plasma, is obtained after centrifugation of whole blood in the presence of anti-coagulants (e.g. EDTA, heparin or sodium citrate). This isolation eliminates cellular material and leaves cell-free components available for detailed characterization. In this review, we focus on proteins found in plasma and discuss how to achieve robust data using mass spectrometry (MS)-based approaches.

Plasma proteomics has undergone a revival in the last five years. The need for more clinically-translatable biological insights is driving an increase in the number of MS-based proteomic studies^3^. Currently, there are more than 150 FDA-approved and laboratory developed tests (LDTs) that utilize plasma for protein-based assays, such as C-reactive protein (CRP) levels for coronary disease and insulin levels for diabetes^4^. This existing clinical infrastructure and familiarity with plasma allows for translation of new discoveries from the laboratory into clinical settings^5^.

Plasma is a challenging biological matrix, due to both a large dynamic range in protein expression and the capabilities of state-of-the-art analytical methods. For example, the plasma peptidome^6,7^ or those peptides carried by the human leukocyte antigen (HLA) molecules^8^ represent low abundance, small molecular weight species. Methods other than MS-based techniques, including a variety of high-sensitivity and throughput plasma protein immunoassays^9^ can be used to profile human plasma ^10^. For the circulating antibodies, multiplexed protein or peptide arrays are commonly used to study reactivity towards the autoimmune components of the proteome^11^ and their post-translational modifications^12^. Today there are > 1000 autoantigens found across a large variety of human conditions ^13^. Numerous studies have focused on plasma processing workflows towards achieving a more comprehensive characterization of the plasma proteome^3,14^; some described in the Plasma Processing Workflows section below. While the basic research community might focus on improving depth of coverage to detect low abundance plasma proteins (i.e. ≤1 ng/mL), the clinical community emphasizes reproducibility of measurements and low coefficients of variation to support actionable clinical decisions. In the former, considerations for translating findings from the laboratory to the clinic can be limited as added sample processing steps may create hurdles fort the wide adoption of a new method. In the latter, while the focus may be on high abundance proteins (i.e. >1 ug/mL)^4^ that have evidence for some clinical indication (e.g. CRP and insulin), these may not be sufficiently sensitive for applications such as early disease detection. We suggest there may be a balanced approach that can satisfy the needs of the entire plasma proteomics and clinical research communities?

### Can MS-based plasma proteomics overcome current challenges?

A key challenge in human health, and an unmet need of medicine, is early disease detection, which is almost entirely dependent on more specific biomarkers, better patient stratification, and methods for predicting patient response to treatment. MS-based plasma proteomics can deliver solutions to many of these challenges when applied in an appropriate manner. Today, there is strong protein-level evidence (i.e., PE1 data) for 17,694 proteins of the human proteome^15^. In a recent 2017 update, HUPO’s Human Plasma Proteome Project (HPPP) reported 3,500 detectable proteins in plasma, emerging from almost 180 studies with a protein-level FDR of 1%^16^. This represents about 20% of the whole currently detectable human proteome. This number nearly doubled from the 1,929 proteins reported in 2011^17,18^ and points to significant improvements made in analytical sensitivity. However, challenges remain due to the low consistency in reproducible observations among plasma proteomics studies. In fact, in data from ∼ 180 studies collected from 2005-17, the 500 most abundant plasma proteins were only reported in 50% of studies based on a reanalysis of all datasets by PeptideAtlas with a global protein-level FDR applied on the ensemble of all 180 studies (Figure 1), hence missing data and coverage of protein abundance across all study subjects will be important aspects of plasma proteomic studies. While some of these differences can be explained by use of different instruments, plasma processing techniques and sample collection methods, it also raises additional questions about which factors most strongly influence plasma protein detectability? Towards answering this question, and improving the overall performance and coverage of plasma proteomics experiments, we recommend that researchers consider key components of a proposed plasma proteomics workflow outlined in Figure 2 below.

**Figure 1.**
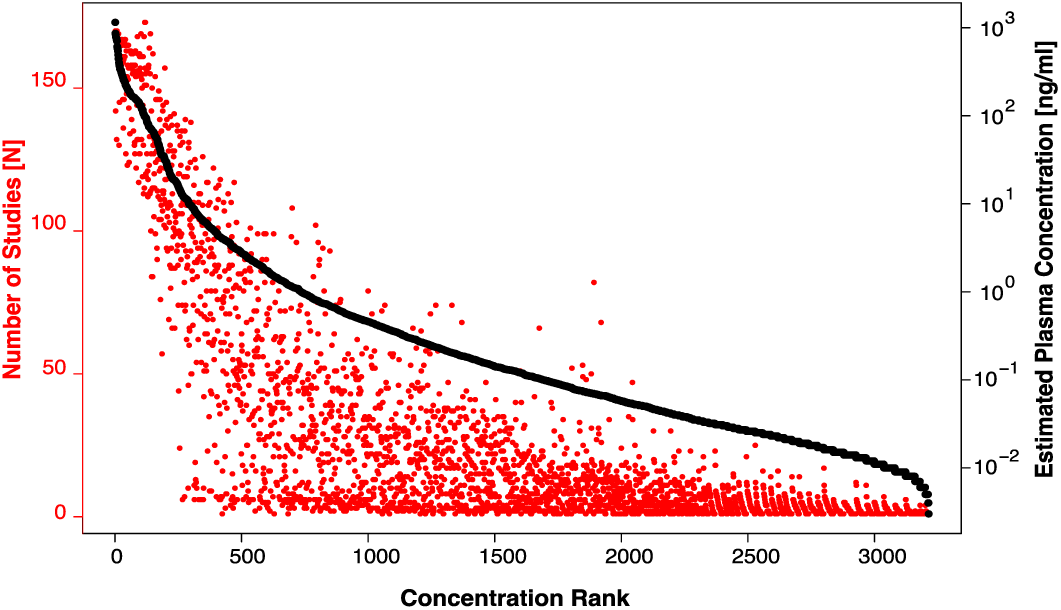
Frequency of protein identification in relation to plasma and serum concentration.

**Figure 2.**
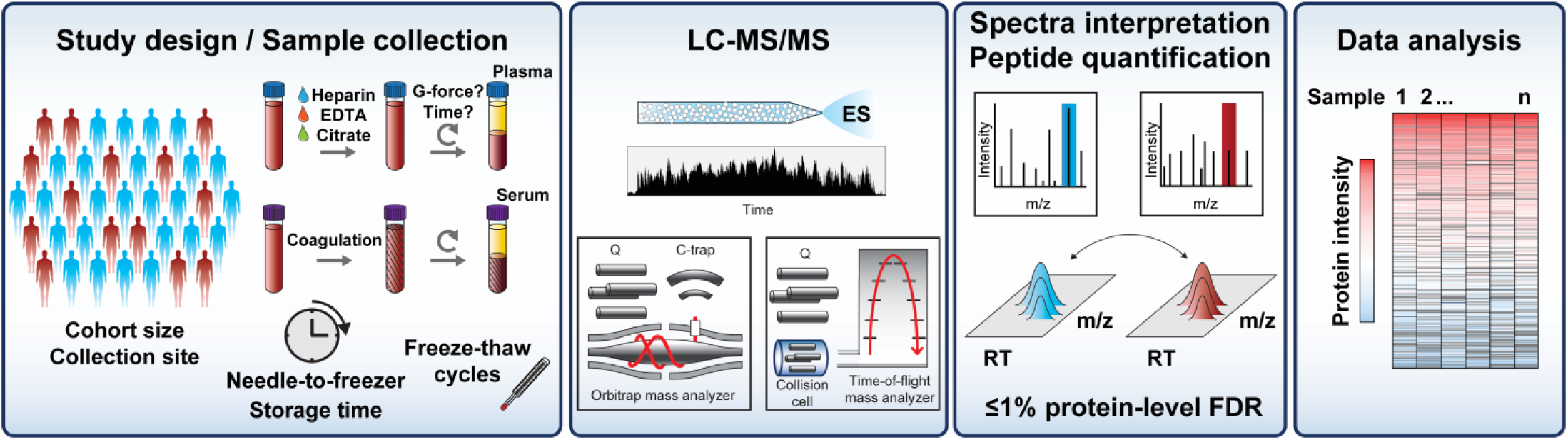
Components of a plasma proteomics workflow. Profiling proteins in plasma begins with collecting the samples in a standardized manner, reporting pre-analytical variables related to the sample and information about the blood donors. After protein digestion and peptide purification, the peptides are separated by liquid chromatography (LC) and ionized by electro spray (ES) for the analysis in the mass spectrometer (MS). Appropriate MS workflows and peptide identification and quantification tools are then applied. For protein identifications, the HPP guidelines recommend a protein-level FDR of ≤1%. Lastly, data analysis should consider how many peptides and proteins were identified and their consistency across samples.

Using data collected for the 2017 draft of the human plasma and serum proteome (hosted by PeptideAtlas^16^), proteins (in red) are plotted as a function of their concentration rank (x-axis) and the number of studies in which they were identified (y-axis, left). The identified proteins (solid black line) are also plotted as a function of their concentration rank (x-axis) and their estimated concentration (y-axis, right). The data are compiled from 178 samples, 71% of which are from plasma, 14% from serum, and 15% of unclear origin.

### Components of a plasma proteomics study

There are numerous considerations to take into account when planning and executing a plasma proteomics study. Some of these are more general whilst others are specific to the analysis of this particular human biospecimen type. In this review, we focus on the elements that span a full project, namely; (1) study design, (2) plasma collection and processing, (3) data acquisition by MS, (4) peptide identification and quantification, and (5) bioinformatic and statistical analysis. As summarized in Figure 2, we introduce these different layers of information across these areas. Lastly, we discuss how each one can impact the outcome of a particular plasma proteomics study.

## Study design

A well-designed plasma proteomics study requires a clear research question along with *a priori* hypotheses that defines the purpose of any study and data analysis. The proteins identified by such studies might be used to compare different stages of a disease or measure the effect of treatment in a given set of patients. By defining and deciding on study-specific factors early, subsequent study protocols and experimental design decisions can be made appropriately. Furthermore, this deliberate approach will ensure that data processing and bioinformatic analysis are executed in a purposeful manner. This also ensures that if subsequent analysis is performed (e.g. after reviewing the preliminary results), this can be correctly categorized as *post-hoc* analysis. Here, we discuss study design considerations that can be grouped into the following categories: study settings, cohort selection, and reference samples.

### Study settings (e.g. specific disease, healthy, or drug investigation)

Plasma proteomics studies to date, and especially in the last five years, have converged in three areas: (1) techniques to improve proteome coverage (i.e. credibly detect the largest number of plasma proteins), (2) solutions that are applicable for clinical applications (e.g. sample throughput, reproducibility, and costs), and (3) studies investigating diverse diseases (e.g. cardiovascular diseases, cancers) or the effect of therapeutics on the plasma proteome (e.g. chemotherapy).

### Cohort selection - sample size

Historically, plasma proteomic studies have small sample sizes - typically <100. This can be attributed to difficulties in sourcing plasma samples with sufficient quality, particularly high sample processing costs (e.g. depletion and/or fractionation) and limitations in data acquisition throughput. More recently, efforts to generate large sample biobanks for proteomic analysis ^19,20^, the introduction of automated and high-throughput sample preparation workflows^21–23^, and improvements in liquid chromatography have facilitated larger cohort studies^23^. Some developments combine rapid sample preparation protocols, multiplexing strategies, automated platforms and optimized HPLC setups^21,24–26^. Beyond these technical aspects, there is a growing recognition that separating biological signal from sample variability often requires large sample cohorts. Accordingly, in an ideal situation, sample size itself would not be a limiting factor during the study design process. Impressively, this has allowed researchers to measure the proteome in cohorts of hundreds to thousands of samples^27–35^.

While large sample sizes can facilitate better powered proteomic studies, they introduce additional experimental considerations aimed at avoiding the introduction of bias into data analysis. In particular, large sample numbers result in an increased data acquisition time, either on one or across multiple instruments. Appropriate design of technical and experimental considerations is required to group samples into processing batches in a balanced and randomized manner, minimizing introduction of bias that could result from acquisition time, run order, operator and/or instrument. Typically, a combination of instrument performance, sample-related variables (e.g. age of sample, inclusion order, time point of collection), and donor-related metadata (e.g. age, sex, ethnicity, disease state) are used to set the maximum number of samples within a processing batch, and the randomization of samples across those batches. When executed optimally, large-scale studies will shift research from small-scale discovery phase to the so-called “rectangular” study designs, where large sample numbers can be analyzed in both discovery and validation stages of biomarker research ^3^. In this way, large cohort studies could enable a significant paradigm shift in the utility of plasma proteomics for clinical applications.

### Cohort selection - age (adult vs. pediatric)

According to the 2019 Revision of World Population Prospects, 25% of the world’s population will be under 15 years of age in 2020, 16% between 15 and 25, 50% between 25 and 64 years of age, and 10% above 65 years of age ^36^. Despite this distribution, an often-underappreciated aspect of previous plasma proteomics studies is that a majority of studies have focused on adults, with only a small proportion of published studies targeting children (approximately 0.6%). Researchers should keep the population age distribution in mind when selecting samples for any study. This is especially important when considering early disease detection is critical for children, especially when trying to limit both short- and long-term sequelae of any disease. Additionally, a recent proteogenomic study revealed that newborns have three times the number of unique proteins as their mothers, further suggesting that differences in plasma proteomes between adults and children could lead to novel biological outcomes ^37^. Studies focusing on sick children are critical for understanding underlying population-specific pathophysiologies and may help reduce guesswork in medical interventions currently associated with drug dosage and predicted patient response. A recent review by McCafferty *et al*., summarizes 35 plasma proteomic studies focused on biomarker discovery in pediatric populations ^35^.

### Cohort selection - reference samples

Selection of reference samples is an important, study-specific consideration for all plasma proteomic projects, as this typically forms the basis for any comparative data analysis. For example, consider a study attempting to identify plasma biomarkers of mild traumatic brain injury (mTBI) to identify subjects who will have delayed outcomes. In this case, the ideal cohort consists of pre-trauma samples from all subjects to assess individual baseline values, the mTBI subjects who have delayed outcomes, and reference samples consisting of mTBI subjects who do not have delayed outcomes that have been matched for age, sex and ethnicity. While this approach seems appropriate, it is important to recognize that epidemiological studies over the last several decades have revealed that there are sometimes additional variables to consider when matching study groups, such as medication, disease history, or state of hydration. Matching of these variables will reduce the impact of bias in data, and hence stratified randomization becomes a feasible solution. However, some of these variables could remain unknown during planning phases of a study (denoted here as “hidden variables”). Accounting for hidden variables is challenging, and sometimes the only course of action is with a sample size large enough that any study will be robust against these potential effects. Depending on the contribution of experimental batches on generated data, total randomization would be preferable over a stratified randomization.

In addition to reference samples for matching conditions/groups, the use of “healthy/normal” subjects as a reference group also needs to be made cautiously. First, a subject included in a “healthy/normal” group may not be representative of the general population and/or the intended use population (in the case of clinical tests), as their selection and inclusion may be based on practical rather than clinically useful criteria. Secondarily, if the “healthy/normal” group is only tested to be negative for a particular disease/condition (e.g. mTBI without delayed outcomes) they could still be positive for other conditions. Consequently, the categorization as “diagnosis-free” may be a more appropriate stratification, as this implies that other diseases or conditions may exist. One approach to mitigate against such risk in comparative analysis between cases and controls is to compare within groups of cases (e.g. mild versus severe disease), rather than those that would otherwise have no condition. For example, a study investigating biomarkers of aging may not need a specific negative control group. Instead, study samples could be representative of the age spectrum in question and ensure that other patient variables (e.g. sex and ethnicity) are balanced across those age groups.

### Reference ranges

In addition to selecting the correct set of reference samples, understanding the range of expected values for reference samples is equally critical during planning plasma proteomics studies. For example, the field of developmental proteomics has demonstrated that biological systems (e.g., hemostatic and inflammatory) in healthy populations undergo age-specific changes in protein expression, from neonates to adults ^38,39^. This is particularly important in diagnostic testing and biomarker discovery applications, where adequate information about the “healthy/normal” groups should be collected as a continuum from birth until adulthood. Moreover, plasma protein expression levels can be individual-specific, where they are stable over time within an individual but vary considerably between individuals ^31,32^. To control for this variability, it may be necessary to define individual-specific or “personalized” reference values rather than population level values ^3,4^. This approach occurs in some longitudinal multi-omics studies ^40^, where subjects are followed temporally and are effectively self-controlled, helping distinguish whether differences between study groups are due to phenotype of interest or other traits (e.g. medication, genetics, or lifestyle).

## Plasma generation and quality metrics

Pre-analytical decisions can have a large impact on the quality and consistency of plasma proteomic data. It is therefore recommended that researchers obtain a clear understanding of how plasma samples were/should be, collected, processed, and stored. The importance in differentiating between quality (e.g. defined by degree of hemolysis) and integrity (e.g. defined by capability to detect a protein of interest in biobanked samples) plays a crucial role in plasma proteomics study design.

During blood collection, factors that need to be considered or monitored include phlebotomy procedures (e.g. needle gauge, number of times subject is exposed to blood sampling) and blood collection tube type (e.g. vacuum container, coagulation activators or inhibitors). Likewise, during plasma processing, factors that can affect the quality of the prepared samples include centrifugation speed, duration, braking rate, temperature, delay from blood collection, plasma processing and storage and where samples were collected. The fact that blood as a biological fluid experiences a change in temperature from 37°C to 20°C during venous draw, may introduce additional and not fully understood physiological processes should not be overlooked. Apart from plasma storage conditions, the number of freeze-thaw cycles add further pre-analytical variables^41^; the relative abundance of S-cysteinylated albumin can provide an estimate of the time for which specimens have been exposed to thawed conditions^42^. In particular, extensive delays of several hours prior to separating blood cells from fluids can alter plasma proteome composition due to erythrocyte and platelet degradation^43^. Protein degradation may further occur due to instability or the action of proteases or depletion of active inhibitors (e.g. SERPINs). At the same time, other studies suggest that a large variety of plasma proteins are relatively stable to post-centrifugation delay and number of freeze-thaw cycles^44,45^ which may indicate that sedimented cells or the use of gel plugs to keep cells apart from fluids may be beneficial for protein detectability. Nevertheless, it is important to establish standard operating procedures (SOPs) and strictly follow these in order to minimize systematic pre-analytical variation that otherwise can result in significant plasma proteome change.

In case retrospective samples are being accessed, and the opportunity for defining protocols controlling for the pre-analytical process is not possible, alternative strategies can be used to help minimize systematic bias within a study. In “freezer studies”, where samples are selected from global biobanks, it may be necessary to evaluate preparatory quality and integrity of available plasma samples. First, centrifugation protocols for plasma generation should be reviewed because these directly influence abundance and types of cellular material that could be present in plasma. Specifically, single-spun plasma typically contains many platelets, and these samples could be affected by post-collection release of platelet proteins. Alternatively, double-spun centrifugation protocols have the capacity to remove a significant number of contaminating cells, including platelets. In order to minimize impact of post-collection platelet activation, double-spun plasma should be used wherever possible. Second, recently generated reference proteomes for erythrocytes and platelets and comparisons of plasma with serum resulted in three “contamination panels” against which one should benchmark samples^46^. There, the high prevalence of cellular proteins might suggest contamination and samples can be flagged for further investigation.

Despite all efforts to control for pre-analytical variability, some degree of systematic bias is inevitable when working with human samples. However, such bias can be minimized through appropriate study design decisions to create randomized and appropriately balanced processing batches. Further, information about pre-analytical variables concerning the sample collection (e.g. needle-to-freezer time, timepoint of sample collection, center or geographical location of sampling) should be collected as a common procedure and considered as part of the data analysis and quality assurance process. Related issues with systematic bias are already known and controlled for in other proteomics applications, including large-scale affinity proteomics experiments^47,48^. In Table 1, we present examples with observations of how sample related pre-analytical variables have affected proteomics data to date.

**Table 1:**
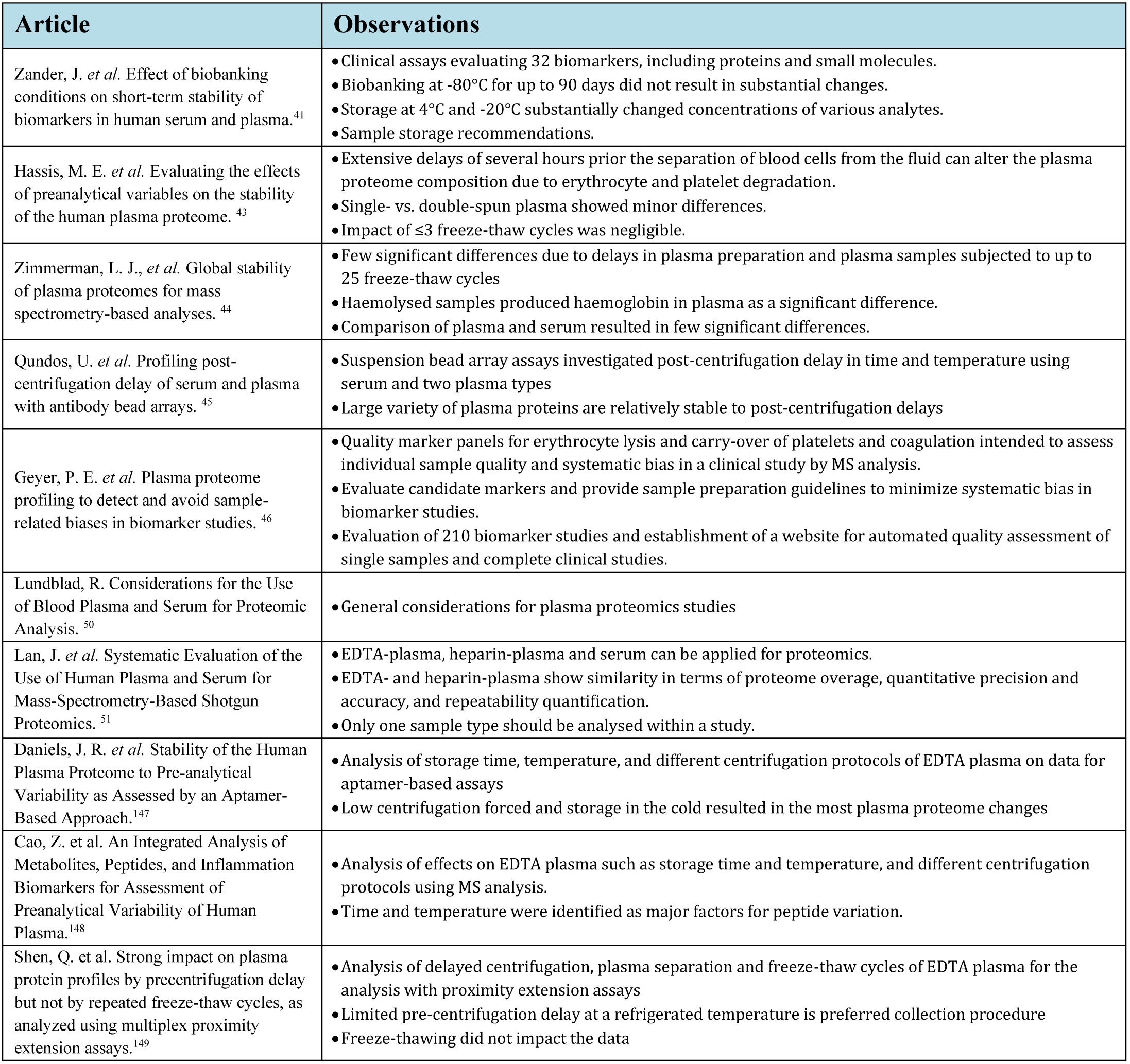
Examples of publications evaluation the influence of pre-analytical sample processing and storage on the sample integrity and preparatory quality of the plasma proteome.

### Plasma is not equivalent to serum

The distinction between plasma and serum as different biological sample types is important when it comes to executing a proteomics study on blood-derived products^49^. For example, designing the most appropriate study plan, correctly interpreting results and appropriately comparing and contrasting results require knowledge of sample type. To clarify this distinction: plasma is obtained by centrifugation of whole blood, while serum is obtained after blood clotting and centrifugation. By removing the blood clot during the preparation of serum, some high abundance proteins such as fibrinogen will be drastically decreased in their concentration in serum but will be present in plasma, increasing the ability to detect some low abundance proteins in serum. At the same time, there are many proteins that are either actively involved in clot formation, nonspecifically adsorbed to clotting proteins, or randomly captured during clot formation^46,50^. For example, the process of whole blood coagulation induces protein secretion from platelets, amongst other cell types; inconsistencies in this process between samples can lead to false positives for differentially expressed proteins in serum. At the same time, clot formation and/or removal processes could be impacted by phenotype (e.g., disease or age), and therefore such differences could represent real biological signal.

While the differences between plasma and serum are largely appreciated in clinical practice, these are underestimated by the proteomics community. Occasionally, the terms serum and plasma are even used interchangeably. However, understanding the differences between these two sample types is critical for advancing proteomics towards the clinic. Systematic comparison of plasma and serum by MS-based proteomics and affinity-based assays point out the many clear differences in composition of these sample types ^51,52^. In 2005, the first recommendation made by the HPP’s HPPP suggested EDTA plasmas be the preferred sample type for all proteomics experiments^49^. Indeed, > 70% of all data sets collected for the plasma proteome draft in 2017^16^ were generated with plasma, however, serum still is the preferred samples type for testing biochemical analytes in clinical chemistry^53^.

## Plasma processing workflows

Comprehensive characterization of the plasma proteome is difficult. The large dynamic range of circulating proteins, combined with the diversity of known and unknown protein isoforms, complicates any analysis, including liquid chromatography coupled to MS (LC-MS)^54^. However, motivated by the high value of identifying and characterizing plasma proteins, standard proteomics workflows have expanded to include prefractionation and labeling approaches to address such challenges^55–57^. Here, we provide a brief overview of these pre-analytical methods and highlight some recent examples.

### Depletion workflows

Although the abundance of plasma proteins spans a large dynamic range, which can be greater than 12 orders of magnitude. Albumin accounts for 50% and the most abundant 22 proteins account for 99% of plasma proteins by weight^54^. This characteristic can be utilized to improve detectability of low abundance proteins by systematic depletion of high abundance proteins^58^. For example, immuno-depletion spin columns, immuno-depletion-LC and magnetic beads have been commercialized to remove up to the 20 most abundant plasma proteins^59–64^. This strategy has also been extended to deplete moderately abundant proteins (e.g., complement proteins, fibronectin, plasminogen) using Affibody molecules, bead-bound peptide hexamers, or antibodies^65 60,66,67^. In addition to depletion strategies using affinity reagents, methods utilizing nanoparticles also exist^61,68,69^. These nanoparticles are designed to distinguish proteins based on physical properties and can be tuned to both exclude (i.e., deplete albumin) or enrich (i.e., capture small proteins^70^ or specific analytes) simultaneously. While depletion methods can help access lower abundance plasma proteins, using these reagents has limitations, including increased sample handling, lower reproducibility/throughput and carry-over concerns (i.e., when depleting multiple samples consecutively)^61^. Moreover, it is important to recognize that many proteins are bound to albumin (e.g., the so-called albuminome) and co-depletion can occur as many abundant proteins have carrier functions. Removing these bound passenger proteins may result in less than controllable off-target depletion^66,67^.

### Fractionation workflows

An alternative strategy to immuno- or affinity-based depletion for more comprehensive plasma protein detection is sample fractionation. This can be used alone and/or in combination with immuno- or affinity-based depletion. Although depletion methods could be considered a form of fractionation (i.e., separating the high abundance from the low abundance components), in such workflows the “high abundance” component is treated as waste. Here, fractionation is defined as a method that divides plasma proteins into usable groups for subsequent characterization. In one approach, two-dimensional liquid chromatography (2D-LC) can divide one plasma sample into many, less complex fractions, whereby the standard low pH reversed-phase (RP) LC conditions applied in most LC-MS workflows is coupled with an orthogonal LC method. While early implementations of 2D-LC used strong cation exchange chromatography prior to standard RPLC methods (e.g., multidimensional protein identification technology, or MudPIT)^71^, there has recently been a shift towards high pH reversed-phase chromatography^72,73^. In both cases, the specific number of fractions can vary, as well as how those fractions are combined, if at all, prior to LC-MS analysis^74^. Additionally, these methods can be executed online with a mass spectrometer, or off-line where samples are first fractionated and fractions are then separately subjected to LC-MS. Fractionation can also be achieved by gel electrophoresis, including in-gel digestion or as a desalting mechanism, where unwanted salts/materials are removed^75^. Recently, a high-resolution isoelectric focusing approach has been developed and utilized for MS-based plasma analysis^37^. Regardless of implementation, one general concern with fractionation-based methods is the increase in the number of samples that needs to be measured, as this scales with the chosen number of fractions. Consequently, fractionation methods create challenges with regards to sample throughput and normalizing data across multiple LC-MS runs.

### Enrichment workflows

In cases where comprehensive characterization of the entire plasma proteome is not necessary, a strategy around enrichment and concentration might be better suited. In this approach, workflows are used to increase the sensitivity for specific proteins of interest^62^. For example, methods have been developed to focus on specific sub-proteomes (plasma glycoproteome and phosphoproteome)^76–78^, proteins that have a specific activity (cytokines for signaling), or those that originate from specific compartments (membrane proteins)^79^. In addition to functional-based enrichment, physical properties of proteins have also been utilized to increase target-protein concentrations^69,80^. For example, proteins can be selectively precipitated by salts or organic solvents^81^; separated by size using chromatography, dialysis, membrane filtration concentrators, or gel electrophoresis^80,82,83^; and further separated by charge (e.g., abundant glycosylation). Finally, single or multiplexed enrichment is possible using immunoaffinity reagents (e.g., antibodies, aptamers, or derivatives thereof), peptides, or chemical baits^84–86^, and they can be used to identify individual proteins or groups of interacting proteins in plasma^87^.

### Quantification workflows

Quantification (relative and/or absolute) is a critical aspect of nearly all plasma proteomics studies. The relative abundance of proteins is typically measured by label-free or isobaric labeling techniques. The most common classes of reagents for isobaric labeling are the tandem mass tags (TMTs)^88^ and the isobaric tag for relative and absolute quantitation (iTRAQ)^89,90^. These tags permit multiplexing of several samples per LC-MS run, commonly 4, 6, 8, or 10 samples, but more are possible^91^. To learn more about the use of isobaric labeling for plasma proteomics, see review by Moulder et al.,^92^ and recent applications^37,93,94^. Multiplexing methods compensate for throughput concerns associated with large sample numbers and some plasma processing workflows (e.g., fractionation), enabling the combination of several samples^95,96^. At the same time, isobaric-based labeling reagents have drawbacks, including lowering target sensitivity (i.e., the signal from individual samples can be diluted below the detection limit if samples have low signal) and reduction in quantitation accuracy (i.e., co-isolated peptides interfere with quantitation due to ratio distortion known as “ratio compression”). In the latter, there are options available to minimize the impact of ratio compression through software correction, narrow isolation windows, further fragmentation of the peptide fragment ions and novel isobaric tags^97–99^. Relative protein abundance can also be measured using label-free techniques. While this approach does not require additional reagents or sample processing, it does not allow sample multiplexing.

In contrast, absolute abundance of specific target proteins can be determined by spiking heavy-labeled reference peptides or proteins at a known concentration. Similarly to sample stability considerations, these reference materials should undergo their own quality control process to ensure assumptions about concentrations or content are accurate. When used correctly, such spike-ins are an attractive concept to serve as a reference between studies, instruments and laboratories. The reference can derive from peptides or proteins and can be added prior to or after protein digestion. Utility of these concepts has recently been demonstrated by selecting suitable standards from a large library of protein fragments^100^, or by spiking-in peptides using stable isotope standards and capture by anti-peptide antibodies (SISCAPA)^101^.

### General workflow considerations

With any sample processing workflow, characterizing its technical performance is as important as the potential value that the method may afford. All too often, one method may work well in the hands of one researcher but cannot be reproduced by another researcher. This has had the unfortunate consequence of leading to opposing or orthogonal workflows for similar goals, making it harder for untrained experts to make informed decisions when implementing plasma proteomics workflows in their own labs. The purpose of applying a particular workflow may be justified for one application (e.g. depletion in order to build a peptide library), while it can be less suitable for others (e.g. profile large number of samples). However, the consequences of applying different enrichment strategies using more recent MS methods and peptide identification guidelines have just recently been investigated^102^. The study showed that about 210 plasma proteins (42% of all the identified) were common among different peptide fractionation methods and 180 proteins (41% of all identified) could be found in either depleted or non-depleted plasma.

Additionally, the lack of technical performance information has led to assays with low reproducibility, resulting in hard-to-replicate results. The community should therefore consider to work towards simplified workflows because each additional sample preparation or processing step, such as depletion, can introduce uncertainty, variance or bias into the data. These concerns are particularly relevant for plasma proteomics studies where large-cohort projects are typically required to make biological conclusions, and minimizing technical noise is critical in identifying small, biologically significant changes. Towards this goal, researchers should consider incorporating broader technical performance characterization into their method development process, and should understand features such as multi-laboratory repeatability and identify common sources of confounding in their method of choice ^103,104^. This information can help the plasma proteomics community move towards the ultimate goal of improved plasma characterization for applications in human health and biology.

## Data acquisition by mass spectrometry

There are broadly two approaches to measure peptides in plasma via MS: targeted and untargeted workflows.

### Targeted plasma proteomics

Selected reaction monitoring (SRM) is the typical targeted approach, wherein target peptides must be selected in advance and the instrument programmed with the expected signatures of those peptides, thus enabling the measurement of relative ion abundances or upper limits for each of the desired targets in every sample^105^. Such an approach does require a potentially time-intensive, up-front process of initial target selection and signature transition optimization, although resources such as SRMAtlas ^106^ enable rapid selection of target peptides and their signatures. Targeting of peptides in plasma can be quite challenging since plasma is a very high dynamic range and complex background in which the target peptides must undergo careful validation procedures to be confirmed and quantified, as demonstrated by the SpecTRA study group^107^. A popular strategy relies on the use of spiked-in stable isotope standards (SIS) as reference peptides to help ensure the correct molecules are being identified^108^. Also, Carr et al. have proposed an important set of guidelines, organized by three tiers of rigor, for the application of SRM to biological samples^109^. In general, targeted proteomics is the method of choice when the number of analytes is relatively small and known in advance, and quantitative measurements are a crucial requirement of the experiment.

### Untargeted plasma proteomics

For untargeted proteomics, there are two broad approaches: data-dependent acquisition (DDA) and data-independent acquisition (DIA). In DDA workflows, the mass spectrometer acquires survey scans to assess which precursor ions are currently entering the instrument, and then sequentially selects several of them to isolate and fragment in turn ^110^. The fragmentation spectrum ideally contains the fragments of just a single precursor. In the DIA workflow (such as with SWATH-MS^111^), the instrument usually also acquires survey scans every few seconds, but then in between scans it steps through a series of selection windows, often 25 m/z units wide, producing fragmentation spectra of all precursors in the wide window multiplexed together ^112^. The advantage of the DIA workflow is that the fragmentation patterns of all precursor ions within the selected mass ranges are recorded, unlike for DDA, wherein fragmentation data is only collected for selected precursors in a semi-stochastic manner. The disadvantage of the DIA workflow is that more complex and less mature software is required to demultiplex the very dense fragmentation spectra. However, the substantial advantage is that with a successful analysis there are typically far fewer missing values in the final data matrix. An additional substantial difference is that while DDA workflows are amenable to label-free quantitation as well as isotopic and isobaric labelling, DIA workflows typically rely on label-free quantitation. As DIA acquires only signals of a distinct m/z-window at a given time, peptides from very high abundant proteins are less of a problem in general^111^. The same benefit for plasma samples has been demonstrated for the DDA method BoxCar, in which multiple m/z-windows are filled to increase the ion injection time compared to a single full MS scan^113^. The potential of both approaches has recently been demonstrated for plasma analysis using isobaric labeled DDA^96^ and in studies using DIA^28,30–32,114^.

## Data processing and bioinformatic analysis

The data generated by mass spectrometers is generally quite complex and requires substantial downstream analysis with sophisticated software tools^115^. However, as far as these software tools are concerned, analysis of data sets derived from plasma samples does not differ substantially from that of other sample types, such as tissue or urine. For SRM data analysis, in addition to vendor-provided tools, Skyline^116^ dominates the free and open-source software field. For DDA data analysis, there are many analysis tools available, including MASCOT^117^, SEQUEST^118^, MaxQuant^119^, and X!Tandem^120^, just to name a few; please see a recent review by Nesvizhskii et al.^121^ for a more comprehensive list. For DIA data analysis, the options are far fewer than for DDA, with OpenSWATH^122^, Spectronaut^123^, PeakView, and DIA-Umpire^124^ as the most frequently used tools. For most SRM and DIA analysis workflows, the tools for identification and quantitation are integrated and work together by default. For many DDA data analysis workflows, the identification and quantitation components are separate tools and the compatibility of those tools is important.

The proteomics community has already done a great job of lowering barriers and working towards freely accessible data in public databases. This approach to openness should continue. It is now common to deposit proteomics datasets in data repositories, most of which are members of the ProteomeXchange Consortium^125,126^. ProteomeXchange sets basic standards and minimum requirements for its members and fosters similar submission and dissemination policies. The main repositories of ProteomeXchange are PRIDE^127^, PeptideAtlas^128,129^ (with its SRM component PASSEL^130^), MassIVE^131^, jPOST^132^, iProX^133^, and Panorama Public^134^. Researchers are encouraged not only to deposit their final datasets in a ProteomeXchange repository, but also to consider downloading and examining previously downloaded and generated datasets to inform the generation of their own data.

There are various formal guidelines that should be followed when submitting manuscripts describing a plasma analysis depending on the type of data and publication. Some journals have their own specific sets of guidelines, such as for the Journal of Proteome Research and Molecular and Cellular Proteomics^135^. Contributions as part of the Human Proteome Project (HPP) must follow the HPP MS Data Interpretation Guidelines ^136^. Other guidelines are applicable to certain workflows, such as for DIA data^137^ and targeted SRM data^109^. It is well worth preparing for the relevant guidelines in advance of data analysis, since complying with some guidelines after an analysis is complete may require redoing some of the work.

## Conclusion

Here we summarize several key aspects about performing plasma proteomics experiments using MS. We provide insights into current capabilities but also raise awareness about the challenges that remain to be addressed. This complements other reviews on MS-based plasma proteomics and its route towards greater translational utility^3,14^. From the perspective of “How should you perform your plasma proteomics experiment?”, we discuss several design elements that are often omitted when focusing on improving the technology rather than its application. While there is not a single “correct” way of performing plasma proteomics, comparative analysis of different methods, such as those proposed for antibody validation^138^, would be a valuable path forward. In addition, we suggest that considering the list of 1000 “popular” plasma proteins that can be detected by MS and affinity-based methods^16^ might be a useful approach.

Given that there are different endpoints and aims for conducting plasma proteomics assays, it remains difficult to suggest a single way of choosing the components of the plasma proteomics pipeline and to determine which criteria should be applied to assess the quality of a preparation or the sample integrity. While pre-analytical information about retrospective samples already stored in biobanks cannot always be retrieved, some caution should be applied when generalizing about the future utility of the observed outcomes. For newly established studies, it will be essential to collect as much information about the samples as possible and determine the influence of pre-analytical variables on the experimental data. Similar to other guidelines, the community could, for example, agree to provide pre-analytical metadata with the deposition of the experimental data, such as the collection site, dates of sample collection, storage times and the Sample PREanalytical Code (SPREC)^139^. As a long-term effort, a framework for storing even donor related metadata alongside pre-analytical and experimental data should be discussed.

In an era with growing numbers of multi-omics studies, we also suggest to consider the sample types and preparations preferred by other methods. These methods may provide additional constrains and reasons for choosing one sample type and collection protocol over the other. As described for metabolomics^140–142^, a common suggestion is to process samples as quickly as possible (e.g. < 30 min) and store these soon as possible at −80°C. We also recommend to use one sample type per study, and if limited availability requires to switch to another preparation type, then there is a need to compare sample types that have been collected and prepared at one blood draw.

Another aspect will be to increase the number of proteins commonly identified in plasma by MS. This could indeed include to collectively annotate which the expected plasma proteins are and determine their susceptibility for pre-analytical variables. This should then also include the available standards, reference peptides or proteins. In addition to the provided sample related criteria, we suggest to consider the impact of post translational modifications on the data. Here, differential glycosylation pattern induced genetic variants have been shown to enable a classification of individual donors^143^. Besides the technological advances, different protocols and studied disease phenotypes, genetic variance that induces a variance in the protein sequences could indeed be another viable reason why only 50% of all MS studies detected 500 common proteins. Hence, data analysis pipelines should evaluate common sequence variance in case genetic data from the blood donor is not available^144^. An example of how sequence variance of other genes than the one of interest can influence protein levels is given by the blood group defining gene *ABO*, which has been shown to affect the plasma levels of von Willebrand factor (VWF)^145^.

Ultimately, the usefulness of proteomics as a methodological approach is dependent on its clinical applicability as a tool to improve patient outcomes. Unfortunately, plasma proteomic studies have, for the most part, focused more on identifying the largest number of proteins rather than focusing on the proteins that can be detected consistently and that have a clinical utility (e.g., predicting a clinical outcome). This raises a couple of questions: Are we, as proteomics community, cooperating enough towards the common goal on translating plasma proteomics across research labs and into the clinics? Are we aware of the issues faced in the clinical setting and do we understand how proteomics can assist ^146^? Is this the reason why proteomics is not as advanced as genomics or transcriptomics when it comes to translational research and clinical utility? Considering the technological advances in proteomics that are actionable from a clinical perspective is certainly a key component in getting proteomics into the clinic, and doing so more quickly and effectively. There is no question that plasma proteomics can have a clear and significant impact on improving clinical diagnostics.

The future of plasma proteomics in the context of diagnostic laboratories is highly reliant on knowledge of the normal, age-specific expression ranges for plasma proteins and their use for accurate diagnosis for our population as a whole. With advanced research in this field, plasma proteomics can provide a reliable, efficient, and highly capable approach to take proteomics to the clinic, to drive a truly personalized medicine experience, and, most importantly, to contribute to human health. While capitalizing from the rapid advances in mass spectrometry, a greater diversity of data from well-designed biomarker discovery and validation studies will become available. Hence, plasma proteomics is well on the way to developing a robust set tools for quantifying proteins across major diseases that will be translated into robust assays made available to diagnostic laboratories.

## Notes

Krishnan K. Palaniappan is an employee of Freenome. All other authors declare no competing financial interest.

## Acknowledgements

We thank the community for the providing open access data. We thank Conor McCafferty for assistance with this manuscript. This work was funded in part by the National Institutes of Health grants R01GM087221 (EWD), R24GM127667 (EWD), U19AG02312 (EWD), U54ES017885 (GSO), and U24CA210967-01 (GSO). JMS acknowledges support from The KTH Center for Applied Precision Medicine, funded by the Erling-Persson Family Foundation grant for Science for Life Laboratory, and the Knut and Alice Wallenberg Foundation. PEG acknowledges support from the Max Planck Society for the Advancement of Science and by the Novo Nordisk Foundation (grant NNF15CC0001).

## References

1. Jin, X. et al. Serum biomarkers of colorectal cancer with AU and NP20 chips including a diagnosis model. Hepatogastroenterology 59, 124–129 (2012).

2. Taguchi, A. & Hanash, S. M. Unleashing the power of proteomics to develop blood-based cancer markers. Clin. Chem. 59, 119–126 (2013).

3. Geyer, P. E., Holdt, L. M., Teupser, D. & Mann, M. Revisiting biomarker discovery by plasma proteomics. Mol. Syst. Biol. 13, 942 (2017).

4. Anderson, N. L. The clinical plasma proteome: a survey of clinical assays for proteins in plasma and serum. Clin. Chem. 56, 177–185 (2010).

5. Percy, A. J. et al. Clinical translation of MS-based, quantitative plasma proteomics: status, challenges, requirements, and potential. Expert Rev Proteomics 13, 673–684 (2016).

6. Dufresne, J. et al. The plasma peptidome. Clin Proteomics 15, 39 (2018).

7. Parker, B. L. et al. Multiplexed Temporal Quantification of the Exercise-regulated Plasma Peptidome. Mol. Cell Proteomics 16, 2055–2068 (2017).

8. Bassani-Sternberg, M. et al. Soluble plasma HLA peptidome as a potential source for cancer biomarkers. Proc. Natl. Acad. Sci. U.S.A. 107, 18769–18776 (2010).

9. Chen, Z., Dodig-Crnkovic, T., Schwenk, J. M. & Tao, S.-C. Current applications of antibody microarrays. Clin Proteomics 15, 7 (2018).

10. Smith, J. G. & Gerszten, R. E. Emerging Affinity-Based Proteomic Technologies for Large-Scale Plasma Profiling in Cardiovascular Disease. Circulation 135, 1651–1664 (2017).

11. Ayoglu, B., Schwenk, J. M. & Nilsson, P. Antigen arrays for profiling autoantibody repertoires. Bioanalysis 8, 1105–1126 (2016).

12. Atak, A. et al. Protein microarray applications: Autoantibody detection and posttranslational modification. Proteomics 16, 2557–2569 (2016).

13. Wang, D. et al. AAgAtlas 1.0: a human autoantigen database. Nucleic Acids Res. 45, D769–D776 (2017).

14. Pernemalm, M. & Lehtiö, J. Mass spectrometry-based plasma proteomics: state of the art and future outlook. Expert Rev Proteomics 11, 431–448 (2014).

15. Omenn, G. S. et al. Progress on Identifying and Characterizing the Human Proteome: 2019 Metrics from the HUPO Human Proteome Project. J. Proteome Res. (2019). doi:10.1021/acs.jproteome.9b00434

16. Schwenk, J. M. et al. The Human Plasma Proteome Draft of 2017: Building on the Human Plasma PeptideAtlas from Mass Spectrometry and Complementary Assays. J. Proteome Res. 16, 4299–4310 (2017).

17. Farrah, T. et al. The State of the Human Proteome in 2012 as Viewed through PeptideAtlas. Journal of Proteome Research 12, 162–171 (2013).

18. Farrah, T. et al. A high-confidence human plasma proteome reference set with estimated concentrations in PeptideAtlas. Mol. Cell Proteomics 10, M110.006353 (2011).

19. Malm, J. et al. Developments in biobanking workflow standardization providing sample integrity and stability. J Proteomics 95, 38–45 (2013).

20. Glimelius, B. et al. U-CAN: a prospective longitudinal collection of biomaterials and clinical information from adult cancer patients in Sweden. Acta Oncol 57, 187–194 (2018).

21. Geyer, P. E. et al. Plasma Proteome Profiling to Assess Human Health and Disease. Cell Systems 2, 185–195 (2016).

22. Fu, Q. et al. Highly Reproducible Automated Proteomics Sample Preparation Workflow for Quantitative Mass Spectrometry. J. Proteome Res. 17, 420–428 (2018).

23. Yu, Y., Bekele, S. & Pieper, R. Quick 96FASP for high throughput quantitative proteome analysis. J Proteomics 166, 1–7 (2017).

24. Bache, N. et al. A Novel LC System Embeds Analytes in Pre-formed Gradients for Rapid, Ultra-robust Proteomics. Mol Cell Proteomics 17, 2284–2296 (2018).

25. Kulak, N. A., Pichler, G., Paron, I., Nagaraj, N. & Mann, M. Minimal, encapsulated proteomic-sample processing applied to copy-number estimation in eukaryotic cells. Nature Methods 11, 319–324 (2014).

26. Dayon, L., Núñez Galindo, A., Cominetti, O., Corthésy, J. & Kussmann, M. A Highly Automated Shotgun Proteomic Workflow: Clinical Scale and Robustness for Biomarker Discovery in Blood. Methods Mol. Biol. 1619, 433–449 (2017).

27. Yin, X. et al. Plasma Proteomics for Epidemiology: Increasing Throughput With Standard-Flow Rates. Circ Cardiovasc Genet 10, e001808 (2017).

28. Wewer Albrechtsen, N. J. et al. Plasma Proteome Profiling Reveals Dynamics of Inflammatory and Lipid Homeostasis Markers after Roux-En-Y Gastric Bypass Surgery. Cell Syst 7, 601-612.e3 (2018).

29. Lee, S. E. et al. Plasma Proteome Biomarkers of Inflammation in School Aged Children in Nepal. PLoS ONE 10, e0144279 (2015).

30. Bruderer, R. et al. Analysis of 1508 Plasma Samples by Capillary-Flow Data-Independent Acquisition Profiles Proteomics of Weight Loss and Maintenance. Mol. Cell Proteomics 18, 1242–1254 (2019).

31. Geyer, P. E. et al. Proteomics reveals the effects of sustained weight loss on the human plasma proteome. Molecular Systems Biology 12, 901 (2016).

32. Liu, Y. et al. Quantitative variability of 342 plasma proteins in a human twin population. Mol. Syst. Biol. 11, 786 (2015).

33. Cominetti, O. et al. Proteomic Biomarker Discovery in 1000 Human Plasma Samples with Mass Spectrometry. J. Proteome Res. 15, 389–399 (2016).

34. Cominetti, O. et al. Obesity shows preserved plasma proteome in large independent clinical cohorts. Sci Rep 8, 16981 (2018).

35. McCafferty, C., Chaaban, J. & Ignjatovic, V. Plasma proteomics and the paediatric patient. Expert Review of Proteomics 16, 401–411 (2019).

36. World Population Prospects 2019: Highlights | Multimedia Library-United Nations Department of Economic and Social Affairs. Available at: https://www.un.org/development/desa/publications/world-population-prospects-2019-highlights.html. (Accessed: 3rd July 2019)

37. Pernemalm, M. et al. In-depth human plasma proteome analysis captures tissue proteins and transfer of protein variants across the placenta. Elife 8, (2019).

38. Ignjatovic, V. et al. Age-related differences in plasma proteins: how plasma proteins change from neonates to adults. PLoS One 6, e17213 (2011).

39. Bjelosevic, S. et al. Quantitative Age-specific Variability of Plasma Proteins in Healthy Neonates, Children and Adults. Molecular & Cellular Proteomics: MCP 16, 924–935 (2017).

40. Price, N. D. et al. A wellness study of 108 individuals using personal, dense, dynamic data clouds. Nat. Biotechnol. 35, 747–756 (2017).

41. Zander, J. et al. Effect of biobanking conditions on short-term stability of biomarkers in human serum and plasma. Clinical Chemistry and Laboratory Medicine 52, 629–639 (2014).

42. Jeffs, J. W. et al. Delta-S-Cys-Albumin: A Lab Test that Quantifies Cumulative Exposure of Archived Human Blood Plasma and Serum Samples to Thawed Conditions. Mol. Cell Proteomics (2019). doi:10.1074/mcp.TIR119.001659

43. Hassis, M. E. et al. Evaluating the effects of preanalytical variables on the stability of the human plasma proteome. Anal Biochem 478, 14–22 (2015).

44. Zimmerman, L. J., Li, M., Yarbrough, W. G., Slebos, R. J. C. & Liebler, D. C. Global stability of plasma proteomes for mass spectrometry-based analyses. Mol. Cell Proteomics 11, M111.014340 (2012).

45. Qundos, U. et al. Profiling post-centrifugation delay of serum and plasma with antibody bead arrays. J Proteomics 95, 46–54 (2013).

46. Geyer, P. E. et al. Plasma proteome profiling to detect and avoid sample-related biases in biomarker studies. bioRxiv 478305 (2018). doi:10.1101/478305

47. Candia, J. et al. Assessment of Variability in the SOMAscan Assay. Sci Rep 7, 14248 (2017).

48. Hong, M.-G., Lee, W., Nilsson, P., Pawitan, Y. & Schwenk, J. M. Multidimensional Normalization to Minimize Plate Effects of Suspension Bead Array Data. J. Proteome Res. 15, 3473–3480 (2016).

49. Omenn, G. S. et al. Overview of the HUPO Plasma Proteome Project: results from the pilot phase with 35 collaborating laboratories and multiple analytical groups, generating a core dataset of 3020 proteins and a publicly-available database. Proteomics 5, 3226–3245 (2005).

50. Lundblad, R. Considerations for the Use of Blood Plasma and Serum for Proteomic Analysis. The Internet Journal of Genomics and Proteomics 1, 8 (2003).

51. Lan, J. et al. Systematic Evaluation of the Use of Human Plasma and Serum for Mass-Spectrometry-Based Shotgun Proteomics. J. Proteome Res. 17, 1426–1435 (2018).

52. Schwenk, J. M. et al. Comparative protein profiling of serum and plasma using an antibody suspension bead array approach. Proteomics 10, 532–540 (2010).

53. Hoq, M. et al. Reference Values for 30 Common Biochemistry Analytes Across 5 Different Analyzers in Neonates and Children 30 Days to 18 Years of Age. Clin. Chem. (2019). doi:10.1373/clinchem.2019.306431

54. Anderson, N. L. & Anderson, N. G. The human plasma proteome: history, character, and diagnostic prospects. Mol. Cell Proteomics 1, 845–867 (2002).

55. Geyer, P. E., Holdt, L. M., Teupser, D. & Mann, M. Revisiting biomarker discovery by plasma proteomics. Mol. Syst. Biol. 13, 942 (2017).

56. Mesmin, C., Oostrum, J. van & Domon, B. Complexity reduction of clinical samples for routine mass spectrometric analysis. PROTEOMICS – Clinical Applications 10, 315–322 (2016).

57. Zhang, Q., Faca, V. & Hanash, S. Mining the Plasma Proteome for Disease Applications Across Seven Logs of Protein Abundance. J. Proteome Res. 10, 46–50 (2011).

58. Gianazza, E., Miller, I., Palazzolo, L., Parravicini, C. & Eberini, I. With or without you - Proteomics with or without major plasma/serum proteins. J Proteomics 140, 62–80 (2016).

59. Tan, S.-H., Mohamedali, A., Kapur, A. & Baker, M. S. Ultradepletion of human plasma using chicken antibodies: a proof of concept study. J. Proteome Res. 12, 2399–2413 (2013).

60. Beer, L. A., Ky, B., Barnhart, K. T. & Speicher, D. W. In-Depth, Reproducible Analysis of Human Plasma Using IgY 14 and SuperMix Immunodepletion. Methods Mol. Biol. 1619, 81–101 (2017).

61. Tu, C. et al. Depletion of abundant plasma proteins and limitations of plasma proteomics. J. Proteome Res. 9, 4982–4991 (2010).

62. Wu, C., Duan, J., Liu, T., Smith, R. D. & Qian, W.-J. Contributions of immunoaffinity chromatography to deep proteome profiling of human biofluids. J. Chromatogr. B Analyt. Technol. Biomed. Life Sci. 1021, 57–68 (2016).

63. Hakimi, A., Auluck, J., Jones, G. D. D., Ng, L. L. & Jones, D. J. L. Assessment of reproducibility in depletion and enrichment workflows for plasma proteomics using label-free quantitative data-independent LC-MS. Proteomics 14, 4–13 (2014).

64. Pringels, L., Broeckx, V., Boonen, K., Landuyt, B. & Schoofs, L. Abundant plasma protein depletion using ammonium sulfate precipitation and Protein A affinity chromatography. J. Chromatogr. B Analyt. Technol. Biomed. Life Sci. 1089, 43–59 (2018).

65. Eriksson, C., Schwenk, J. M., Sjöberg, A. & Hober, S. Affibody molecule-mediated depletion of HSA and IgG using different buffer compositions: a 15 min protocol for parallel processing of 1-48 samples. Biotechnol. Appl. Biochem. 56, 49–57 (2010).

66. Shi, T. et al. IgY14 and SuperMix immunoaffinity separations coupled with liquid chromatography-mass spectrometry for human plasma proteomics biomarker discovery. Methods 56, 246–253 (2012).

67. Thulasiraman, V. et al. Reduction of the concentration difference of proteins in biological liquids using a library of combinatorial ligands. Electrophoresis 26, 3561–3571 (2005).

68. Gökay, Ö., Karakoç, V., Andaç, M., Türkmen, D. & Denizli, A. Dye-attached magnetic poly(hydroxyethyl methacrylate) nanospheres for albumin depletion from human plasma. Artif Cells Nanomed Biotechnol 43, 62–70 (2015).

69. Capriotti, A. L. et al. Comparison of three different enrichment strategies for serum low molecular weight protein identification using shotgun proteomics approach. Anal. Chim. Acta 740, 58–65 (2012).

70. Harney, D. et al. Small-protein enrichment assay enables the rapid, unbiased analysis of over 100 low abundance factors from human plasma. Mol. Cell Proteomics (2019). doi:10.1074/mcp.TIR119.001562

71. Washburn, M. P., Wolters, D. & Yates, J. R. Large-scale analysis of the yeast proteome by multidimensional protein identification technology. Nat. Biotechnol. 19, 242–247 (2001).

72. Dwivedi, R. C. et al. Practical implementation of 2D HPLC scheme with accurate peptide retention prediction in both dimensions for highthroughput bottom-up proteomics. Anal. Chem. 80, 7036–7042 (2008).

73. Delmotte, N., Lasaosa, M., Tholey, A., Heinzle, E. & Huber, C. G. Two-dimensional reversed-phase x ion-pair reversed-phase HPLC: an alternative approach to high-resolution peptide separation for shotgun proteome analysis. J. Proteome Res. 6, 4363–4373 (2007).

74. Cao, Z., Tang, H.-Y., Wang, H., Liu, Q. & Speicher, D. W. Systematic comparison of fractionation methods for in-depth analysis of plasma proteomes. J. Proteome Res. 11, 3090–3100 (2012).

75. Abdallah, C., Dumas-Gaudot, E., Renaut, J. & Sergeant, K. Gel-based and gel-free quantitative proteomics approaches at a glance. Int J Plant Genomics 2012, 494572 (2012).

76. Fanayan, S., Hincapie, M. & Hancock, W. S. Using lectins to harvest the plasma/serum glycoproteome. Electrophoresis 33, 1746–1754 (2012).

77. Hendriks, I. A. et al. Site-specific mapping of the human SUMO proteome reveals co-modification with phosphorylation. Nat. Struct. Mol. Biol. 24, 325–336 (2017).

78. Dufresne, J. et al. The plasma peptides of ovarian cancer. Clin Proteomics 15, 41 (2018).

79. Glisovic-Aplenc, T. et al. Improved surfaceome coverage with a labelfree nonaffinity-purified workflow. Proteomics 17, (2017).

80. Kim, B. et al. Affinity enrichment for mass spectrometry: improving the yield of low abundance biomarkers. Expert Rev Proteomics 15, 353–366 (2018).

81. Henning, A.-K., Albrecht, D., Riedel, K., Mettenleiter, T. C. & Karger, A. An alternative method for serum protein depletion/enrichment by precipitation at mildly acidic pH values and low ionic strength. Proteomics 15, 1935–1940 (2015).

82. De Bock, M. et al. Comparison of three methods for fractionation and enrichment of low molecular weight proteins for SELDI-TOF-MS differential analysis. Talanta 82, 245–254 (2010).

83. Onnerfjord, P., Eremin, S. A., Emnéus, J. & Marko-Varga, G. High sample throughput flow immunoassay utilising restricted access columns for the separation of bound and free label. J Chromatogr A 800, 219–230 (1998).

84. Razavi, M. et al. High-throughput SISCAPA quantitation of peptides from human plasma digests by ultrafast, liquid chromatography-free mass spectrometry. J. Proteome Res. 11, 5642–5649 (2012).

85. Ippoliti, P. J. et al. Automated Microchromatography Enables Multiplexing of Immunoaffinity Enrichment of Peptides to Greater than 150 for Targeted MS-Based Assays. Anal. Chem. 88, 7548–7555 (2016).

86. Devine, M., Juba, M., Russo, P. & Bishop, B. Structurally stable N-t-butylacrylamide hydrogel particles for the capture of peptides. Colloids Surf B Biointerfaces 161, 471–479 (2018).

87. Fredolini, C. et al. Systematic assessment of antibody selectivity in plasma based on a resource of enrichment profiles. Sci Rep 9, 8324 (2019).

88. Thompson, A. et al. Tandem mass tags: a novel quantification strategy for comparative analysis of complex protein mixtures by MS/MS. Anal. Chem. 75, 1895–1904 (2003).

89. Ross, P. L. et al. Multiplexed protein quantitation in Saccharomyces cerevisiae using amine-reactive isobaric tagging reagents. Mol. Cell Proteomics 3, 1154–1169 (2004).

90. Pottiez, G., Wiederin, J., Fox, H. S. & Ciborowski, P. Comparison of 4-plex to 8-plex iTRAQ quantitative measurements of proteins in human plasma samples. J. Proteome Res. 11, 3774–3781 (2012).

91. McAlister, G. C. et al. Increasing the multiplexing capacity of TMTs using reporter ion isotopologues with isobaric masses. Anal. Chem. 84, 7469–7478 (2012).

92. Moulder, R., Bhosale, S. D., Goodlett, D. R. & Lahesmaa, R. Analysis of the plasma proteome using iTRAQ and TMT-based Isobaric labeling. Mass Spectrom Rev 37, 583–606 (2018).

93. Dey, K. K. et al. Deep undepleted human serum proteome profiling toward biomarker discovery for Alzheimer’s disease. Clin Proteomics 16, 16 (2019).

94. Du, X. et al. Alterations of Human Plasma Proteome Profile on Adaptation to High-Altitude Hypobaric Hypoxia. J. Proteome Res. 18, 2021–2031 (2019).

95. Keshishian, H. et al. Quantitative, multiplexed workflow for deep analysis of human blood plasma and biomarker discovery by mass spectrometry. Nat Protoc 12, 1683–1701 (2017).

96. Keshishian, H. et al. Multiplexed, Quantitative Workflow for Sensitive Biomarker Discovery in Plasma Yields Novel Candidates for Early Myocardial Injury. Mol. Cell Proteomics 14, 2375–2393 (2015).

97. Karp, N. A. et al. Addressing Accuracy and Precision Issues in iTRAQ Quantitation. Mol Cell Proteomics 9, 1885–1897 (2010).

98. Virreira Winter, S. et al. EASI-tag enables accurate multiplexed and interference-free MS2-based proteome quantification. Nat Methods 15, 527–530 (2018).

99. Ting, L., Rad, R., Gygi, S. P. & Haas, W. MS3 eliminates ratio distortion in isobaric labeling-based multiplexed quantitative proteomics. Nat Methods 8, 937–940 (2011).

100. Edfors, F. et al. Screening a Resource of Recombinant Protein Fragments for Targeted Proteomics. J. Proteome Res. 18, 2706–2718 (2019).

101. Anderson, N. L., Razavi, M., Pope, M. E., Yip, R. & Pearson, T. W. Multiplexed measurement of protein biomarkers in high-frequency longitudinal dried blood spot (DBS) samples: characterization of inflammatory responses. (Systems Biology, 2019). doi:10.1101/643239

102. Ahn, S. B. et al. Potential early clinical stage colorectal cancer diagnosis using a proteomics blood test panel. Clin Proteomics 16, 34 (2019).

103. Collins, B. C. et al. Multi-laboratory assessment of reproducibility, qualitative and quantitative performance of SWATH-mass spectrometry. Nat Commun 8, 291 (2017).

104. Abbatiello, S. E. et al. Large-Scale Interlaboratory Study to Develop, Analytically Validate and Apply Highly Multiplexed, Quantitative Peptide Assays to Measure Cancer-Relevant Proteins in Plasma. Mol. Cell Proteomics 14, 2357–2374 (2015).

105. Picotti, P. & Aebersold, R. Selected reaction monitoring-based proteomics: workflows, potential, pitfalls and future directions. Nat. Methods 9, 555–566 (2012).

106. Kusebauch, U. et al. Human SRMAtlas: A Resource of Targeted Assays to Quantify the Complete Human Proteome. Cell 166, 766–778 (2016).

107. Penn, A. M. et al. Validation of a proteomic biomarker panel to diagnose minor-stroke and transient ischaemic attack: phase 2 of SpecTRA, a large scale translational study. Biomarkers 23, 793–803 (2018).

108. Kopylov, A. T. et al. 200+ Protein Concentrations in Healthy Human Blood Plasma: Targeted Quantitative SRM SIS Screening of Chromosomes 18, 13, Y, and the Mitochondrial Chromosome Encoded Proteome. J. Proteome Res. 18, 120–129 (2019).

109. Carr, S. A. et al. Targeted peptide measurements in biology and medicine: best practices for mass spectrometry-based assay development using a fit-for-purpose approach. Mol. Cell Proteomics 13, 907–917 (2014).

110. Hein, M. Y., Sharma, K., Cox, J. & Mann, M. Chapter 1 - Proteomic Analysis of Cellular Systems. in Handbook of Systems Biology (eds. Walhout, A. J. M., Vidal, M. & Dekker, J.) 3–25 (Academic Press, 2013). doi:10.1016/B978-0-12-385944-0.00001-0

111. Gillet, L. C. et al. Targeted data extraction of the MS/MS spectra generated by data-independent acquisition: a new concept for consistent and accurate proteome analysis. Mol. Cell Proteomics 11, O111.016717 (2012).

112. Ludwig, C. et al. Data-independent acquisition-based SWATH-MS for quantitative proteomics: a tutorial. Mol. Syst. Biol. 14, e8126 (2018).

113. Meier, F., Geyer, P. E., Virreira Winter, S., Cox, J. & Mann, M. BoxCar acquisition method enables single-shot proteomics at a depth of 10,000 proteins in 100 minutes. Nat. Methods 15, 440–448 (2018).

114. Niu, L. et al. Plasma proteome profiling discovers novel proteins associated with non-alcoholic fatty liver disease. Mol. Syst. Biol. 15, e8793 (2019).

115. Deutsch, E. W., Lam, H. & Aebersold, R. Data analysis and bioinformatics tools for tandem mass spectrometry in proteomics. Physiol. Genomics 33, 18–25 (2008).

116. MacLean, B. et al. Skyline: an open source document editor for creating and analyzing targeted proteomics experiments. Bioinformatics 26, 966–968 (2010).

117. Perkins, D. N., Pappin, D. J., Creasy, D. M. & Cottrell, J. S. Probability-based protein identification by searching sequence databases using mass spectrometry data. Electrophoresis 20, 3551–3567 (1999).

118. Eng, J. K., McCormack, A. L. & Yates, J. R. An approach to correlate tandem mass spectral data of peptides with amino acid sequences in a protein database. Journal of the American Society for Mass Spectrometry 5, 976–989 (1994).

119. Cox, J. & Mann, M. MaxQuant enables high peptide identification rates, individualized p.p.b.-range mass accuracies and proteome-wide protein quantification. Nat Biotechnol 26, 1367–1372 (2008).

120. Craig, R. & Beavis, R. C. TANDEM: matching proteins with tandem mass spectra. Bioinformatics 20, 1466–1467 (2004).

121. Nesvizhskii, A. I. A survey of computational methods and error rate estimation procedures for peptide and protein identification in shotgun proteomics. J Proteomics 73, 2092–2123 (2010).

122. Röst, H. L. et al. OpenSWATH enables automated, targeted analysis of data-independent acquisition MS data. Nat. Biotechnol. 32, 219–223 (2014).

123. Bruderer, R. et al. Extending the Limits of Quantitative Proteome Profiling with Data-Independent Acquisition and Application to Acetaminophen-Treated Three-Dimensional Liver Microtissues. Mol Cell Proteomics 14, 1400–1410 (2015).

124. Tsou, C.-C. et al. DIA-Umpire: comprehensive computational framework for data-independent acquisition proteomics. Nat. Methods 12, 258–264, 7 p following 264 (2015).

125. Vizcaíno, J. A. et al. ProteomeXchange provides globally coordinated proteomics data submission and dissemination. Nat. Biotechnol. 32, 223–226 (2014).

126. Deutsch, E. W. et al. The ProteomeXchange consortium in 2017: supporting the cultural change in proteomics public data deposition. Nucleic Acids Res. 45, D1100–D1106 (2017).

127. Perez-Riverol, Y. et al. The PRIDE database and related tools and resources in 2019: improving support for quantification data. Nucleic Acids Res. 47, D442–D450 (2019).

128. Desiere, F. The PeptideAtlas project. Nucleic Acids Research 34, D655–D658 (2006).

129. Desiere, F. et al. Integration with the human genome of peptide sequences obtained by high-throughput mass spectrometry. Genome Biol. 6, R9 (2005).

130. Farrah, T. et al. PASSEL: the PeptideAtlas SRMexperiment library. Proteomics 12, 1170–1175 (2012).

131. Pullman, B. S., Wertz, J., Carver, J. & Bandeira, N. ProteinExplorer: A Repository-Scale Resource for Exploration of Protein Detection in Public Mass Spectrometry Data Sets. J. Proteome Res. 17, 4227–4234 (2018).

132. Moriya, Y. et al. The jPOST environment: an integrated proteomics data repository and database. Nucleic Acids Res. 47, D1218–D1224 (2019).

133. Ma, J. et al. iProX: an integrated proteome resource. Nucleic Acids Res. 47, D1211–D1217 (2019).

134. Sharma, V. et al. Panorama Public: A Public Repository for Quantitative Data Sets Processed in Skyline. Mol. Cell Proteomics 17, 1239–1244 (2018).

135. Bradshaw, R. A., Burlingame, A. L., Carr, S. & Aebersold, R. Reporting Protein Identification Data: The next Generation of Guidelines. Molecular & Cellular Proteomics 5, 787–788 (2006).

136. Deutsch, E. W. et al. Human Proteome Project Mass Spectrometry Data Interpretation Guidelines 2.1. J. Proteome Res. 15, 3961–3970 (2016).

137. Chalkley, R. J., MacCoss, M. J., Jaffe, J. D. & Röst, H. L. Initial Guidelines for Manuscripts Employing Data-independent Acquisition Mass Spectrometry for Proteomic Analysis. Mol. Cell Proteomics 18, 1–2 (2019).

138. Uhlen, M. et al. A proposal for validation of antibodies. Nat. Methods 13, 823–827 (2016).

139. Lehmann, S. et al. Standard preanalytical coding for biospecimens: review and implementation of the Sample PREanalytical Code (SPREC). Biopreserv Biobank 10, 366–374 (2012).

140. Yin, P., Lehmann, R. & Xu, G. Effects of pre-analytical processes on blood samples used in metabolomics studies. Anal Bioanal Chem 407, 4879–4892 (2015).

141. Yin, P. et al. Preanalytical aspects and sample quality assessment in metabolomics studies of human blood. Clin. Chem. 59, 833–845 (2013).

142. Paglia, G. et al. Influence of collection tubes during quantitative targeted metabolomics studies in human blood samples. Clin. Chim. Acta 486, 320–328 (2018).

143. Lin, Y.-H., Zhu, J., Meijer, S., Franc, V. & Heck, A. J. R. Glycoproteogenomics: A Frequent Gene Polymorphism Affects the Glycosylation Pattern of the Human Serum Fetuin/α-2-HS-Glycoprotein. Mol. Cell Proteomics 18, 1479–1490 (2019).

144. Ting, Y. S. et al. PECAN: library-free peptide detection for data-independent acquisition tandem mass spectrometry data. Nat. Methods 14, 903–908 (2017).

145. Desch, K. C. et al. Linkage analysis identifies a locus for plasma von Willebrand factor undetected by genome-wide association. Proc. Natl. Acad. Sci. U.S.A. 110, 588–593 (2013).

146. Grant, R. P. & Hoofnagle, A. N. From lost in translation to paradise found: enabling protein biomarker method transfer by mass spectrometry. Clin. Chem. 60, 941–944 (2014).

147. Daniels, J. R. et al. Stability of the Human Plasma Proteome to Preanalytical Variability as Assessed by an Aptamer-Based Approach. J. Proteome Res. (2019). doi:10.1021/acs.jproteome.9b00320

148. Cao, Z. et al. An Integrated Analysis of Metabolites, Peptides, and Inflammation Biomarkers for Assessment of Preanalytical Variability of Human Plasma. J. Proteome Res. 18, 2411–2421 (2019).

149. Shen, Q. et al. Strong impact on plasma protein profiles by precentrifugation delay but not by repeated freeze-thaw cycles, as analyzed using multiplex proximity extension assays. Clin. Chem. Lab. Med. 56, 582–594 (2018).

